# A Unique Mouse Model of Early Life Exercise Enables Hippocampal Memory and Synaptic Plasticity

**DOI:** 10.1101/699603

**Authors:** Autumn S. Ivy, Tim Yu, Enikö Kramár, Sonia Parievsky, Fred Sohn, Thao Vu

## Abstract

Aerobic exercise is a powerful modulator of learning and memory. Molecular mechanisms underlying the cognitive benefits of exercise are well documented in adult rodents. Animal models of exercise targeting specific postnatal periods of hippocampal development and plasticity are lacking. Here we characterize a model of early-life exercise (ELE) in male and female mice designed with the goal of identifying critical periods by which exercise may have a lasting impact on hippocampal memory and synaptic plasticity. Mice freely accessed a running wheel during three postnatal periods: the 4^th^ postnatal week (juvenile ELE, P21-27), 6^th^ postnatal week (adolescent ELE, P35-41), or 4^th^-6^th^ postnatal weeks (juvenile-adolescent ELE, P21-41). All exercise groups significantly increased their running distances over time. When exposed to a weak learning stimulus, mice that had exercised during the juvenile period were able to form lasting long-term memory for a hippocampus-dependent spatial memory task. Electrophysiological experiments revealed enhanced long-term potentiation in hippocampal CA1 the juvenile-adolescent ELE group only. Furthermore, basal synaptic transmission was significantly increased in all mice that exercised during the juvenile period. Our results suggest early-life exercise can enable hippocampal memory, synaptic plasticity, and basal synaptic physiology when occurring during postnatal periods of hippocampal maturation.

## Introduction

Exercise during adulthood is highly effective in improving cognitive functions in both humans and rodents^1,2^. In children, clinical studies positively associate higher physical activity levels with improved working memory^3^, academic performance^4^, and structural brain health^5^. Preclinical exercise models using adult rodents have identified underlying neurobiological mechanisms by which exercise may improve hippocampus-dependent memory: exercise augments neurogenesis^6^, vascularization^7^, synapse number^8^, facilitates long-term potentiation (LTP) in DG^9–11^; but see^12^ regarding female mice) and in certain cases, area CA1^13,14^; but see ^15^), and induces a number of synaptic genes and proteins, including the key plasticity-modulating growth factor brain-derived neurotrophic factor (BDNF;^16,17^). There is some evidence that exercise may engage similar plasticity mechanisms in the developing brain^18^; however, studies focusing on the neurobiological sequelae of adolescent exercise are lacking. Yet, it is precisely during early-life development periods that the effects of exercise may have the most long-term benefit on memory and cognitive ability throughout the lifespan^19^.

The cognitive and neurotrophic effects of exercise during adulthood are transient, seeming to fade in days to weeks. For example, one study exercised adult rats for three weeks then performed cognitive testing two-weeks after exercise cessation. These rats had weaker memory performance on a radial arm maze task after being sedentary for two weeks post-exercise. Coupled with this finding were gradual reductions in BDNF mRNA and protein in hippocampus, eventually returning to baseline levels 14 days post-exercise cessation^20^. In contrast, more recent studies focused on early-life exercise find that benefits to spatial memory and hippocampal plasticity persist into adulthood when the exercise is initiated in adolescence^21–23^. This suggests an enduring effect of the adolescent exercise experience on learning and memory^24^. Exact temporal windows by which an early life exercise experience can promote enduring changes in neuronal function, hippocampal plasticity, and behavior have not yet been thoroughly investigated.

The mammalian brain undergoes protracted postnatal development in a region- and functional network-specific manner^25,26^. In humans, neuronal differentiation and synaptogenesis within hippocampus reach adult levels around 3-5 years of age^27^, which is around the time children are able to reliably form long-term spatial memory^28^. In rodents, the hippocampus reaches milestones in synapse density and circuit refinement during the 3^rd^ to 4^th^ postnatal weeks (reviewed in^29,30^) which coincides with development of stable long-term potentiation (LTP^31,32^) and establishment of learning and memory functions^33,34^. A characteristic of postnatal brain maturation is that synaptic connections are over-produced and subsequently pruned^35^, thus rendering these developmental processes exquisitely sensitive to environmental experiences^36^. Processes of postnatal neurodevelopment are likely guided by specific activity- and experience-dependent patterns of gene expression that ultimately inform cell function within circuits^37,38.^ It is thus reasonable to target defined periods of hippocampal structural and functional maturation, using biological mouse models, to uncover temporal windows of postnatal development by which hippocampal function can be persistently modulated by exercise.

In this study we developed a model of voluntary physical activity during specific early life developmental periods in mice (early-life exercise, or ELE) to address the hypothesis that the timing of exercise during postnatal hippocampal maturation can lead to enduring benefits in cognitive function and synaptic plasticity. We will refer to the 4^th^ week of life in a mouse (postnatal days 21-27) as a “juvenile” period, distinct from adolescence (5^th^-6^th^ postnatal weeks, as reviewed in^39,40.^ We show that both male and female mice undergoing exercise during the juvenile period exhibit enabled hippocampus-dependent spatial memory, increased long-term potentiation and changes in basal synaptic physiology. Importantly, our ELE model can be used to uncover temporally specific molecular mechanisms engaged by ELE to influence neuronal function and behavior in a lasting manner.

## Results

### Running behavior and weight gain characteristics of the early-life exercise (ELE) mouse model

Given the developmental timing of voluntary exercise in this model, we postulated that there would be important sex-specific and running group-specific differences in weight gain and distance ran across time. These differences would depend upon the *timing* (juvenile and/or adolescence) and *duration* (1- or 3-weeks) of the early-life exercise exposure. Male and female mice were either weaned directly into cages equipped with a running wheel on P21 (juv ELE and juv-adol ELE groups) or placed into cages without running wheels until P35 (adol ELE group). ELE mice were allowed to run for either one week (juv ELE and adol ELE groups) or three weeks (juv-adol ELE group, Fig. 1A). All mice were pair-housed to avoid the effects of social isolation stress^41^. Mice tended to run during the dark phase. For all running groups, the exercise exposure took place either just before or during adolescent growth spurts in mice (usually occurring during the 4^th^-5^th^ postnatal weeks), thus weight gained between P21 and P42 was compared across sed and ELE groups and between sexes. Two-way ANOVA revealed a significant interaction between sex and ELE group, suggesting that ELE has differential effects on weight gain dependent upon sex (Fig. 1B, *F*_(4,94)_ = 2.48, *p* = 0.0492). *Post hoc* multiple comparisons revealed that in males, juv-adol ELE mice had significantly less weight gain than juv ELE mice (*p* = 0.045). Female mice housed with a stationary wheel had significantly greater weight gain than sed (*p* = 0.0005), juv-adol ELE (*p* = 0.0007), and adol ELE female mice (*p* < 0.0001). These findings suggest that a longer duration of ELE exposure (three weeks) during juvenile / adolescence significantly reduces weight gain in male mice, whereas in female mice, presence of a stationary wheel led to greater weight gain than mice in sedentary cages without running wheel and mice that ran during adolescence.

**Figure 1.**
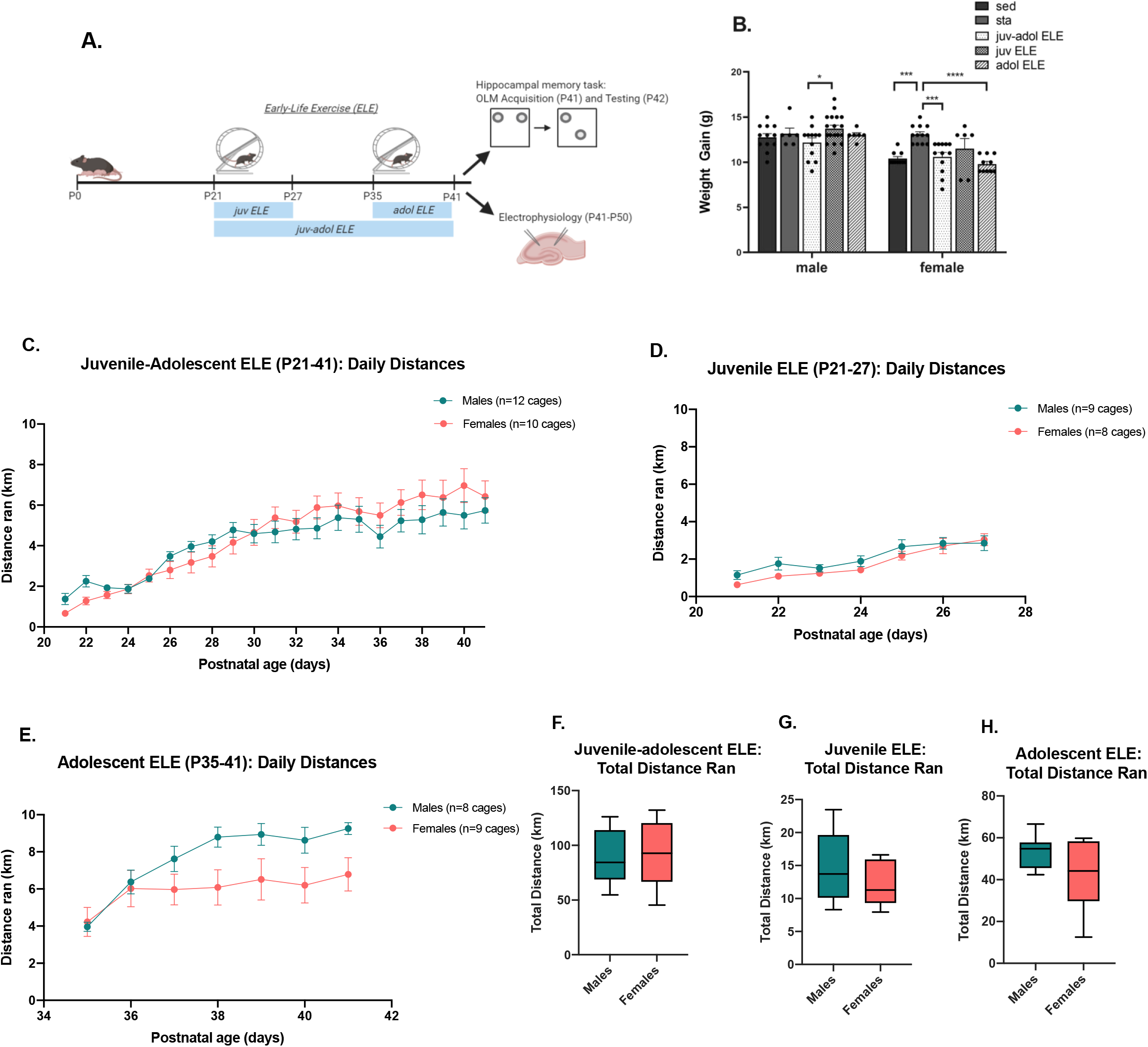
Experimental Design, Running Behavior and Weight Trends in Early Exercise Model. (A) Experimental design diagram. Upon weaning on P21, wild-type male and female mice in the juv ELE and juv-adol ELE were housed in cages equipped with voluntary running wheels for 1-week (juv ELE) or 3-weeks (juv-adol ELE), respectively. Adol ELE mice were housed in cages on P35 and allowed to run freely for 1-week. All mice were then tested in object location memory (OLM) or sacrificed for electrophysiology studies. (B) A significant difference in weight gain was observed between male juv ELE and juv-adol ELE mice. Female stationary mice gained significantly more weight than sedentary, juv-adol ELE, and adol ELE mice. (C-E) All ELE mice significantly increased their running distance throughout the early-life period of running wheel access. There were no significant sex differences in running trends across postnatal days. No difference in running distance by day in juvenile-adolescent ELE mice, and juvenile ELE mice, significant effect of sex and significant interaction in adolescent ELE mice (F-H) No sex differences in total distance ran in any ELE group. Data were analyzed using one-way ANOVA with post-hoc comparisons or Student’s t tests: ** p ≤ 0.01, and *** p ≤ n = 8-12 exercise cages per group, 2 mice per cage

We next addressed the question whether there are sex differences in daily and cumulative running distances in ELE mice. Juv-adol ELE male and female mice gradually increased their daily running distance over the 3-week period (Fig. 1C). A two-way repeated-measures ANOVA revealed a significant effect of postnatal day (*F*_(3, 57)_ = 41.95, *p* < 0.0001) and significant interaction with sex (*F*_(20, 400)_ = 2.146, *p* = 0.0031). There was no independent effect of sex on daily running distance (*F*_(1, 20)_ = 0.1547, *p* = 0.6982), nor was there a significant difference between sexes in total cumulative distance ran (*p* = 0.6982). In evaluating the exercise behavior of mice running for the shorter duration (1-week), we found that postnatal day had a significant effect on daily running distance in juv ELE mice (Fig. 1D, *F*_(3, 51)_ = 35.93, *p* < 0.0001). There was no independent effect of sex on running distance (*F*_(1,15)_ = 1.11, *p* = 0.3087) and no interaction (*F*_(6,90)_ = 1.201, p = 0.3131). In contrast, adol ELE mice had a statistically significant interaction of postnatal day × sex (Fig. 1E, *F*_(6,84)_ = 4.769, *p* = 0.0003). Although female adol ELE mice trended toward less running toward the end of the adolescent running period this did not reach significance (*p* > 0.05). Similar to all other running groups, adol ELE mice also significantly increased their running across postnatal days (*F*_(4, 58)_ = 24.87, *p* < 0.0001). We next compared daily and cumulative running distances in 1-week runners, expecting that running distance would be significantly greater in adol ELE mice when compared to juv ELE mice (regardless of sex) given their body mass differences when entering running cages. There were no sex differences in total running distance within juv and adol ELE groups (Fig. 1F-H); however, cumulative distance traveled over their respective running periods was significantly greater in the adol ELE group when compared to the juv ELE group, and this was true for both sexes (males: *t*_(15)_ = 11.96, *p* < 0.0001; females: *t*_(14)_ = 4.69, *p* = 0.0004). Total distances ran per cage over the duration of the exercise exposures were the following: juv-adol ELE mice; males = 87.8 ± 6.9 km, females = 92.2 ± 9.3 km; juv ELE mice; males = 14.7 ± 1.8 km, females = 12.29 ± 1.2 km; adol ELE mice; males = 53.6 ± 2.8 m, females = 41.8 ± 6.2 km. In sum, all running groups and both sexes of mice gradually and significantly increased their running distances across time. The cumulative amount of voluntary running throughout the running period was dependent on when early life exercise was initiated (in the juvenile period vs in adolescence).

### 1- or 3-weeks of juvenile exercise enables long-term memory in both male and female mice in a lasting manner

Previous studies have demonstrated that in adult mice, a minimum of 2-3 weeks of exercise is required for exercise-induced improvements in long-term memory^42,43^; however, a minimum amount of exercise needed for benefits to learning and memory in juvenile and adolescent mice has not been established. Furthermore, in adult mice, the exercise-induced memory benefits fade by 3 days after exercise ceases^43^. Whether ELE benefits hippocampal memory for a different duration than in adult mice after the cessation of exercise is an unanswered question. We therefore tested the following hypotheses: 1) that a learning stimulus typically insufficient for long-term memory formation in sedentary wild-type mice can become sufficient after ELE (as it does after 2-3 weeks of exercise in adulthood), and 2) the early-life timing of exercise during juvenile and/or adolescent periods will have a lasting impact on the duration of ELE-induced improvements in long-term memory.

Hippocampus-dependent learning and memory was assessed using an Object Location Memory (OLM) task, a non-aversive spatial memory task that relies on the rodent’s innate preference for exploring novel placement of objects^44^. All mice were tested in OLM during adolescence (P42-43). Mice were exposed to two identical objects for either 3-min or 10-min during the OLM acquisition phase, followed by a 5-min OLM retention test performed 24 hours later (Fig. 2A). During OLM testing one of the objects was placed in a novel location and times spent exploring familiar and novel object locations were quantified and expressed as a discrimination index (DI). The 3-min training exposure is a duration previously established as sub-threshold for supporting long-term memory formation in adult mice, whereas the 10-min training exposure is typically sufficient^45^. Our first experiments tested whether 10-min of training during OLM acquisition was also sufficient for long-term memory formation during adolescence. During the 10-min OLM acquisition trials, no object preference was observed in any of the groups, as indicated by average DIs close to 0 (Fig. 2B, males: F_(3, 27)_ = 0.4740, *p* = 0.7030; females: F_(3,27)_ = 1.931, *p* = 0.1494). During OLM testing 24h later, all 10-min trained groups exhibited strong preference for novel-placed objects, as demonstrated by a significant increase in testing DI when compared to acquisition DIs (Fig. 2C, two-way RM-ANOVA comparing OLM acquisition vs OLM testing within groups. Males: main effect of OLM session: F_(1,27)_ = 143.5, *p* < 0.0001; no effect of running group: F_(3,27)_ = 2.064, *p* = 0.1248; and no interaction: F_(3,27)_ = 0.7298, *p* = 0.5432. Females: main effect of OLM session: F_(1,26)_ = 66.4, *p* < 0.0001; no effect of running group: F_(3,26)_ = 0.7795, *p* = 0.5161; and no interaction: F_(3,26)_ = 0.8047, *p* = 0.5026). Running exposure did not significantly improve memory performance during OLM testing, suggesting a possible plateau of learning reached by all groups after being trained for 10-min the day prior (one-way ANOVA, males: F_(3,27)_ = 1.705, *p* = 0.1896; females: F_(3,26)_ = 0.2346, *p* = 0.8714). Further, there were no differences in total OLM testing object exploration times (Fig. 2D, males: F_(3, 27)_ = 0.560, *p* = 0.6460; females: F(3,26) = 1.645, *p* = 0.2032). These data suggest that performance in the OLM task can be used as a robust measure of long-term memory formation in adolescent male and female mice.

**Figure 2.**
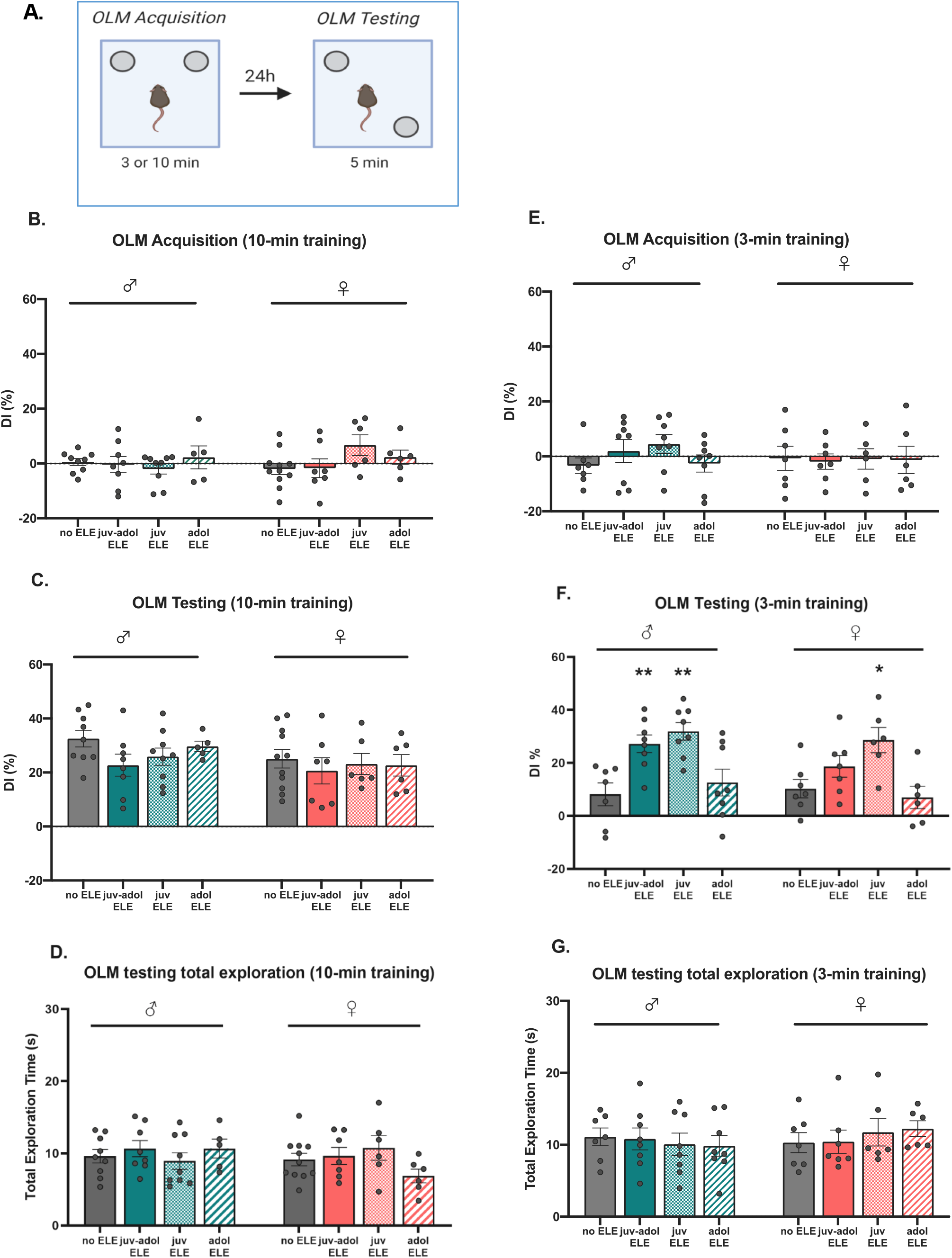
Juvenile-adolescent ELE results in improved novel object recognition memory in male and female mice. (A) Experimental design diagram. (B) OLM acquisition in 10-min trained male and female mice demonstrated no significant object discrimination. (C) During OLM testing, all mice displayed significant exploration of the novel object, and there were no significant differences in DIs between groups. (D) No differences in total exploration during OLM testing in 10-min trained mice. (E) Mice trained for 3-min during OLM acquisition did not demonstrate object preference. (F) Juv ELE and juv-adol ELE male mice had significant preference for the object placed in a novel location, whereas no ELE and adol ELE mice did not. (G) Total exploration times in 3-min trained mice did not differ across groups or sexes. Data were analyzed with *post hoc* Tukey’s comparisons vs ‘no ELE’ mice (* *p* < 0.05; ** *p* < 0.01; *** *p* < 0.005 vs no ELE). N = 6-12 mice per group

3-min of training during OLM acquisition has been demonstrated to be insufficient for long-term memory formation in non-exercised adults but can become sufficient if immediately preceded by exercise^42,43^. Given the exercise-induced improvements in long-term memory in adult mice, we sought to determine whether this phenomenon also holds true during adolescence. All groups trained for 3-min did not exhibit significant object preference during OLM acquisition (Fig. 2E, one-way ANOVA, males: F_(3, 27)_ = 1.174, *p* = 0.3380; females: F_(3,24)_ = 0.0328, *p* = 0.9918). Sedentary male and female mice had significantly lower DIs than 10-min trained sedentary mice, suggesting that 3-min trained, non-exercised adolescent mice do not form strong long-term memory for object location (unpaired *t*-tests, males: t_(14)_ = 4.731, *p* = 0.0003; females: t_(16)_ = 2.904, *p* = 0.0103). Male mice that underwent exercise during the juvenile period (juv ELE and juv-adol ELE; both groups had access to a running wheel during P21-27) showed strong preference for novel object location when trained for 3-min (Fig. 2F, one way ANOVA, F_(3, 27)_ = 7.783, *p* = 0.0007; no ELE vs juv-adol ELE: *p* = 0.0089; no ELE vs juv ELE: *p* = 0.0011) whereas adol ELE mice did not (no ELE vs adol ELE: *p* = 0.8392). In females, only the juv ELE group had significantly greater preference for novel object location (Fig. 2F, one way ANOVA, F_(3, 22)_ = 5.228, *p* = 0.0071; no ELE vs juv ELE: *p* = 0.0142; no ELE vs juv-adol ELE: *p* = 0.3784; no ELE vs adol ELE: *p* = 0.9228). All groups of 3-min trained mice spent similar times exploring objects during OLM testing (Fig. 2G, one-way ANOVA, males: F_(3, 27)_ = 0.1645, *p* = 0.9193; females: F_(3, 22)_ = 0.3922, *p* = 0.7598). These results suggest that in both sexes, 1-week of exercise taking place during a specific juvenile developmental period (P21-27) is sufficient for enhancing long-term memory for an OLM acquisition trial that is usually insufficient for long-term memory in sedentary controls, and furthermore, juv ELE-induced enabling of long-term memory is present at least 2 weeks after the juvenile exercise period ends.

### Synaptic plasticity and basal synaptic physiology in CA1 are modulated in male mice after juvenile exercise

Prior studies have confirmed that memory for object location requires an intact, functioning dorsal hippocampus^46^. Therefore, synaptic plasticity was examined in the CA3-CA1 Schaffer collateral pathway of the hippocampus in sedentary and ELE mice, to test the hypothesis that ELE also leads to an enhancement in synaptic strength in the same hippocampal region likely involved in improved object location memory performance after juv and juv-adol ELE. Long-term potentiation (LTP) studies were performed in acute hippocampal slices from juvenile, adolescent, and juvenile-adolescent ELE male mice. All electrophysiological studies were performed in male mice only, between P42-P51, and induced with a single train of 5 theta bursts to Schaffer collateral inputs and recorded field EPSPs from apical dendrites of CA1b. LTP was notably increased in juv-adol ELE mice but not in the juv ELE or adol ELE groups (Fig 3A, B; one-way ANOVA, F_(3,42)_ = 7.20, p = 0.0005). This is consistent with prior studies demonstrating that one-week of voluntary physical activity is typically insufficient for increased LTP in adult rodents (specifically in dentate gyrus; Farmer et al., 2004). Interestingly, input/output curves from both juv ELE and juv-adol ELE, but not adol ELE mice demonstrated greater level of excitability by revealing stronger fEPSP slopes when compared to sedentary controls, suggesting an up-regulation of basal post-synaptic physiology in both running groups that included the juvenile developmental period (Fig 3C; F_(3,38)_ = 5.93, *p* = 0.002). Importantly, this result demonstrates a lasting effect of juvenile exercise on basal physiology, given that the exercise paradigm for these mice terminated two weeks prior to electrophysiological studies were performed. Paired-pulse facilitation experiments were also performed to interrogate presynaptic function. These demonstrated no difference between groups (Fig 3D; F_(3,41)_ = 1.75, *p* = 0.171). Overall, these results are in contrast to data in adult mice demonstrating lack of enhancement in CA1-LTP after chronic voluntary wheel running^9^; however, they implicate a specific, 1-week period of juvenile exercise that can lead to lasting changes in hippocampal CA1 circuit maturation and plasticity.

**Figure 3.**
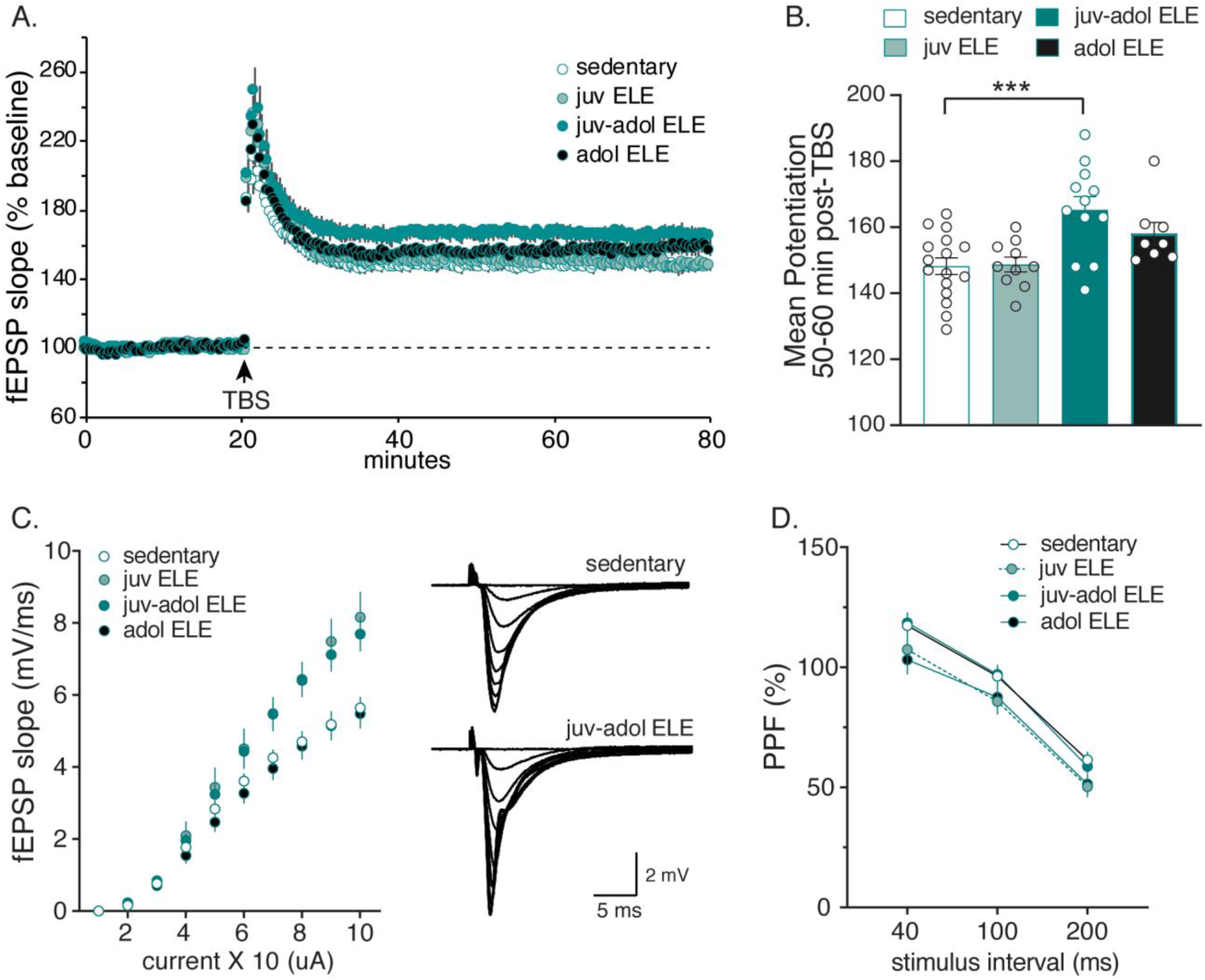
Juvenile exercise leads to enhanced hippocampal synaptic plasticity in CA1. (A) Hippocampal slices from mice undergoing 3 weeks of exercise (juv-adol ELE) had a significant increase in LTP in response to theta burst stimulation (TBS) compared to sedentary controls, 1-week exercise in juvenile mice during P21-27 (juv ELE) and 1-week exercise in adolescent mice during P35-41 (adol ELE). (B) This enhanced potentiation in juv-adol ELE slices maintained significance 50-60 min post-TBS. (C) I/O curves generated in the Schaffer collaterals showed a notable increase in fEPSP slope during the latter part of the curve in both juv ELE and juv-adol ELE compared to adol ELE and sedentary controls. Insert: representative traces collected during generation of I/O curve in slices from sedentary and juv-adol ELE mice. (D) No significant differences were found between groups following paired-pulse facilitation. *** *p* ≤ 0.005

## Discussion

We have successfully designed a mouse model of early-life voluntary exercise for the evaluation of hippocampus-dependent memory and synaptic plasticity during postnatal development. Our goal was to explore the possibility that physical exercise during early life can lead to enduring benefits in hippocampal function. To that end, our model was designed to target early life periods of hippocampal maturation to uncover temporally-specific cellular and molecular mechanisms uniquely engaged by early life exercise. This experimental design allowed us to test the hypothesis that juvenile exercise for one week (P21-27) is sufficient to inform hippocampal function in a lasting manner. Juvenile male and female mice increased their running distance over time with no sex-dependent differences in running activity or weight gained. We discovered that early-life exercise for 4^th^-6^th^ postnatal weeks (juv-adol ELE), as well as a shorter exercise experience only during the 4^th^ postnatal week (juv ELE) both promote hippocampal-dependent spatial memory formation in male mice. This was not true for mice that exercised only during the 6^th^ postnatal week (adol ELE), suggesting that the 4^th^ postnatal week of development was particularly sensitive to the exercise experience with regard to promoting long-term memory in a lasting manner. Extracellular recordings revealed that CA1-LTP was significantly increased only in mice that exercised for 3-weeks but was not further enhanced in the 1-week juvenile, nor 1-week adolescent exercisers when compared to sedentary controls. This finding suggests that one week of voluntary physical activity, regardless of developmental timing, was not sufficient to enhance CA1-LTP in our study. However, a remarkable finding was that basal post-synaptic properties were significantly modulated in both groups of mice that exercised during the 4^th^ postnatal week; furthermore, these changes endured in the juv ELE group, in that input/output studies were performed in this group two-three weeks *after* exercise cessation. This result corresponded with the lasting effect of 1-week juv ELE on enhancing memory function in OLM, which was also demonstrated two weeks after exercise cessation. To our knowledge, this discovery has not been previously tested or described in other existing models of rodent early life or adult exercise.

All mice that exercised during the juvenile period (juv ELE and juv-adol ELE groups) formed long-term memory of a learning event that is typically subthreshold for long-term memory formation (3-min of OLM training;^45^). Most notably, the effect of juv ELE on long-term memory formation persisted in both male and female mice. Our data from the juv-adol ELE group is consistent with prior work demonstrating that in adult male mice, voluntary wheel running for 3-weeks enabled memory in the same hippocampus-dependent OLM paradigm^42^. In that study, administration of the histone deacetylase (HDAC) inhibitor sodium butyrate (NaB) produced effects similar to exercise on memory and maintained memory enhancements for a longer duration than exercise. Further, both exercise and NaB treatment were associated with increased expression of BDNF transcripts I and IV, as well as enriched BDNF promoter acetylation on H4K8 (a marker of transcriptional activation^42,47^). These results suggest that exercise enables memory formation by modifying chromatin structure, and this may permit transcriptional activation of genes required for exercise-induced plasticity and long-term memory formation. Previous work has alluded to this concept as a “molecular memory” of exercise^17,43,48^. Chromatin modifying mechanisms that facilitate the expression of learning- and plasticity-related genes are also engaged by exercise during adolescence. In Abel and Rissman^18^, 1-week of adolescent exercise (P46-52) induced a number of hippocampal genes related to synaptic plasticity and cell signaling (including *Bdnf*, *Cbln1*, *Syn1* and *Syp*), increased global H3 acetylation, and down-regulated a number of HDAC genes^9^. Given the crucial role of histone-modifying mechanisms in memory and synaptic plasticity^49,50^, and our novel finding that juvenile exercise can lead to persistent effects on memory function, a future direction will be to assess whether there is an underlying persistence of chromatin modifications that can increase the efficiency of long-term memory formation after juvenile exercise.

Recent data suggests this is a possibility in adult mice^43^. In their study, mice freely accessed running wheels for specific durations, followed by return to sedentary activity, to determine how long memory enhancements may persist after exercise, and whether they can be reactivated with subsequent short exercise exposure. Key findings from this study were: 1) using the same OLM acquisition paradigm used in this study, they find that a minimum of two weeks of voluntary exercise was required for memory enhancements to persist; 2) these memory enhancements persisted for about three days after return to sedentary activity; and 3) 2wk exercised mice given an additional, 2 day “reactivating exercise” (a duration which by itself is insufficient for enhanced OLM) within a specific temporal window (seven days, but not 14 days, after return to sedentary activity) could re-enhance memory performance in OLM^43^. In our study, one week of exercise was sufficient for memory enhancement in male and female mice, but only when the exercise occurred during the 4^th^ postnatal week. Furthermore, the effects of juvenile exercise on memory enhancement persisted for two weeks after cessation of exercise. These studies both highlight that exercise can have differing impacts on memory performance, dependent on both the duration of exercise exposure, and when in life the exercise took place (during juvenile periods of development or during adulthood). These approaches to studying the effects of exercise on learning and memory can reveal novel neurobiological (potentially epigenetic;^42^) mechanisms underlying the persistence of exercises’ effects on memory, and how they may be differentially engaged in juveniles and adults.

OLM requires an intact dorsal hippocampus^46^. Enabled OLM performance in juv-adol ELE mice complements our finding of enhanced CA1-LTP in these mice. Studies focusing on the effects of exercise on LTP in hippocampal CA1 are relatively underrepresented in the literature since the finding that LTP was unchanged after chronic voluntary wheel running in adult rats^9,10^, but exercise can rescue CA1-LTP impairments in the settings of stress^13^ or sleep deprivation^14^. In our study, however, LTP was enhanced after theta-burst stimulation in the group of animals that underwent three weeks of exercise (juv-adol ELE), but not in those mice that underwent one-week of exercise (juv ELE and adol ELE groups, Fig. 3A). This suggests that during juvenile and adolescent periods, LTP enhancements after early-life exercise may require long-term (greater than 1-week) exercise exposure, which has been suggested previously in adult rodents when LTP is induced in the dentate gyrus^9^. LTP is considered to be the cellular correlate of learning and memory^51^. Theta-burst stimulation was our chosen LTP induction protocol because this form of LTP is dependent on BDNF^52,53^, and potentially mimics circuit firing patterns occurring during learning in rodents^54^ and humans^55^. This study also presents the novel finding that basal synaptic transmission was persistently augmented after juv ELE, as reflected in increased I/O curves from both juv ELE and juv-adol ELE groups. It can be hypothesized that an increased number of functional synapses are available after juv ELE to support increased synaptic transmission, however, this has yet to be explored. In all, further experiments characterizing electrophysiological properties of the hippocampal network after juvenile exercise, particularly when tested well after the exercise experience has ended, would shed light on how synaptic function may be changed in an enduring manner.

With regard to female mice, this study demonstrated that early-life exercise has similar effects on enabling memory in females during juvenile periods, but juv-adol exercise did not lead to improved learning and memory. Prior work has demonstrated that voluntary aerobic physical activity in both male and female rodents during adulthood produces similar benefits in hippocampus-dependent memory tasks (see comprehensive meta-analysis by^56^). Although it has been well established that the neurotrophic factor BDNF is one of the key mediators facilitating the effects of exercise on hippocampal function in male mice^57^, expression of the full-length gene is not similarly enhanced in the hippocampus of female mice after exposure to chronic voluntary wheel running, and there may be sex-dependent variations in BDNF splice-variants after exercise^58^. Also, estrogens have been shown to modulate structural plasticity within hippocampus^59^, can regulate the expression of BDNF, and can enhance hippocampus-dependent memory^60^. In the current study, adolescent females were not formally checked for stage of estrous cycling. C57bl/6J female mice typically enter estrous cycling within 10 days after vaginal opening, which typically occurs around P26^61^. It is very possible that stage of estrous in our ELE mice could have affected overall outcomes in OLM memory enhancement, particularly after juv-adol ELE. Future studies should be designed to time learning and memory testing with stage of estrous cycling; these studies are generally needed to determine whether there exist sex-dependent histone alterations evoked by ELE in influencing hippocampal memory and plasticity genes^62^.

In summary, it is well known that exercise is a highly potent, positive modulator of cognitive function in adults. Yet exercise parameters for optimal cognitive development in typically developing children (as well as cognitively impaired children) currently do not exist. Furthermore, how early life exercise may contribute to later life cognitive functions is an area of research deserving further exploration^19^. Because of the paucity of preclinical and clinical data on the subject, exercise guidelines for children are currently vague^63^. If findings in this study are relevant to humans, they implicate a uniquely lasting effect of exercise during early life on learning and memory functions. Overall, a mechanistic understanding of early-life exercise can be leveraged to inform exercise-based interventions for improving and preserving cognitive functions in children as well as throughout life.

## Materials and Methods

### Animals

Wild-type mice were progeny of C57Bl6J dams, originally obtained from Jackson Laboratories and further bred in our animal facilities. Mice had free access to food and water and lights were maintained on a 12 h light/dark cycle. Upon weaning on postnatal day (P) 21, mice were housed in either standard bedding cages or cages equipped with a running wheel. Both male and female mice were used for all exercise and behavior experiments. Only male mice were used for electrophysiological studies. All behavioral testing was performed during the light cycle. Experiments were conducted according to US National Institutes of Health guidelines for animal care and use and were approved by the Institutional Animal Care and Use Committee of the University of California, Irvine.

### Running Wheel Paradigm

Upon weaning on P21, mice were pair-housed in either standard cages without a wheel, or equipped with either a mobile or stationary stainless-steel running wheel (diameter 4.5 cm, 112 grams, Starr Life Sciences). Animals housed with running wheels had free access for predetermined durations: either one week during the juvenile period (P21-27; juv ELE), one week during adolescence (P35-41; adol ELE), or three consecutive weeks of access (P21-41; juv-adol ELE). Each wheel was fitted with a lightweight polyurethane mesh to prevent the small limbs of weanling mice from slipping through the rungs of the wheels. Running activity was tracked via digitally monitored sensors attached to a data port and computer for real-time data acquisition (Vital View Software) via magnetic detection of the wheel revolutions. Each digitally monitored wheel tracked running distance for the cage (two mice housed per cage, thus distance traveled per cage represents the total distance ran between two mice). Revolutions were quantified by the minute daily for the duration of running and converted to distance ran per cage (km).

### Object Location Memory (OLM) Protocol

The object location memory protocol used in this study has been adapted and modified from previous protocols^44^. On postnatal day 36, mice were brought into a testing room with reduced room brightness. Mice were handled for approximately 2 min each, for a total of 5 consecutive days prior to the OLM training session (twice a day for the first 2 days followed by once a day for the next 3 days). Habituation sessions were 5 min, twice per day for 3 days (P39-P41) and occurred within chambers containing four unique spatial cues on each wall of the chamber (horizontal lines, black X, vertical strip, and blank wall). Habituation overlapped with the last 3 days of handling. During the training phase (P42), mice were placed into the same chambers with two identical objects and exposed to either a subthreshold (3 min) or threshold (10 min) training period; a study design informed by prior work demonstrating that 3 min of training is typically insufficient for long-term memory formation^45^. For mice in exercise cages, running wheels were locked immediately after OLM training to eliminate any acute effects of exercise on memory consolidation. On testing day (P43; 24 hours after training), one of the objects was moved to a novel location inside the chamber, and mice were then allowed to explore the objects for 5 minutes. The times spent with each object was determined via blind analysis by hand-scoring, and values were converted into a Discrimination Index (DI): [(time spent exploring novel object – time spent exploring familiar object)/(total time exploring both objects) × 100%]. There were no significant differences in memory performance between sedentary animals and those housed with a stationary wheel, so these groups were combined in all OLM analyses. Mice with uneven exploration times between the two objects during the training phase (DI > 20) or showed no decrease in total distance traveled across habituation sessions were excluded from analysis.

### Electrophysiology studies

For LTP studies, male mice from all running groups were sacrificed between P42-50, and slices were collected from the rostral hippocampus. Coronal slices were sectioned (Leica VT1000 S) at 320um thickness and placed in an interface recording chamber with preheated (31 ± 1 °C) artificial cerebrospinal fluid (124 mM NaCl, 3 mM KCl, 1.25 mM KH2PO4, 1.5 mM MgSO4, 2.5 mM CaCl2, 26 mM NaHCO3, and 10 mM d-glucose). Slices were continuously perfused at a rate of 1.75–2 ml per min while the surface of the slices was exposed to warm, humidified 95% O2/5% CO2. Recordings began after at least 2 h of incubation. Field excitatory postsynaptic potentials (fEPSPs) were recorded from CA1b stratum radiatum using a single glass pipette (2– 3 MΩ) filled with 2 M NaCl. Stimulation pulses (0.05 Hz) were delivered to Schaffer collateral-commissural projections using a bipolar stimulating electrode (twisted nichrome wire, 65 μm) positioned in CA1c. Current intensity was adjusted to obtain 50% of maximal fEPSP response. After a stable baseline was established, LTP was induced with a single train of five theta bursts, in which each burst (four pulses at 100 Hz) was delivered 200 ms apart (i.e., at theta frequency). The stimulation intensity was not increased during TBS. Data were collected and digitized by NAC 2.0 Neurodata Acquisition System (Theta Burst). Data in LTP figure were normalized to the last 10 min of baseline and presented as mean ± SE. Baseline measures on I/O curves, paired-pulse facilitation and LTP were analyzed using a two-way ANOVA.

### Statistical Analyses

Statistical analyses were performed using either a Student's *t* test, one-way or two-way ANOVAs. Tukey’s *post hoc* tests were used to make specific comparisons when significant interactions, or main effects of sex or running group were observed. For rare instances that 24h running data for a particular day were not recorded, values were imputed based on taking the average distance ran of the prior three days, to perform statistical analyses. Cumulative running distances are expressed as Means ± SD. Two-way repeated-measures ANOVAs were used for the following: 1- to analyze running data and had factors of postnatal day and sex, and 2- to analyze, by each mouse, OLM training vs OLM testing. This latter analysis had factors of OLM session (training vs testing) and ELE group, and *post hoc* comparisons were performed using Sidak’s tests. Main effects and interactions for all ANOVAs are described in the text and figure legends, along with the specific number of animals of each sex used in each experiment. All analyses were two-tailed and required an α value of 0.05 for significance. Error bars in all figures represent SEM. For all experiments, values ±2 standard deviations from the group mean were considered outliers and were removed from analyses. All statistics were performed with GraphPad Prism 8 software. Total number of mice used in behavior experiments = 152.

## Acknowledgements

This research was supported by the National Institute of Neurological Disorders and Stroke of the National Institutes of Health, under award number K12NS098482 (awarded to A.S.I.) and the UC Irvine School of Medicine, Department of Pediatrics. We would also like to thank Dr. Marcelo Wood for scientific discussions and critical input on the project.

